# Model-based pupil deconvolution reveals latent attentional dynamics

**DOI:** 10.64898/2026.09.30.755708

**Authors:** Sangramjit Maity, Sebastiaan Mathôt, Sonia Baloni Ray

**Affiliations:** Department of Experimental Psychology, University of Groningen, The Netherlands; Department of Social Sciences and Humanities, Indraprastha Institute of Information and Technology, New Delhi, India; Centre for Design and New Media, Indraprastha Institute of Information and Technology, New Delhi, India

## Abstract

Changes in pupil size have been widely used as an indirect measure of attention. However, attention engages on a faster timescale than the pupillary response. Therefore, the slow time course of the pupil response makes it difficult to distinguish attentional responses to stimuli presented in close temporal succession. Although signal deconvolution has been used in previous research to estimate latent attentional responses from the pupil signal, existing methods primarily model dilation without considering the visual stimulus evoked constriction phase of the pupil. Here, we introduce a dual-kernel deconvolution model that models constriction and dilation using separate impulse response functions. Using an RSVP paradigm with emotional faces as target stimuli, we demonstrate that the proposed model estimates latent attentional response despite substantial overlap in the measured pupil signal. Importantly, the recovered attentional response resembles the time course of attentional allocation previously predicted by several models of Attentional Blink. Together, these findings demonstrate that the dynamics of latent attentional episodes can be recovered from the pupil response with a temporal resolution previously obscured by the sluggishness of the pupillary response. The proposed method is also provided as a usable software package that serves as a general tool for investigating the temporal dynamics of attention in rapid visual paradigms.

## Introduction

Visual attention is a cognitive mechanism that selectively filters out irrelevant visual information, allowing the brain to focus on task-relevant stimuli. However, it takes time for attention to engage and disengage. Therefore, when multiple targets are presented in close temporal proximity, their successful processing depends on the relative temporal distance between them. The limits of temporal attention are often studied using rapid serial visual presentation (RSVP), in which a series of stimuli is presented in rapid succession at a single location (Chun & Potter, 1995). When two targets are presented in sequence, the second target (T2) is often missed if it appears within 200-500 ms of the first target (T1), a phenomenon known as the Attentional Blink (AB).

The Attentional Blink (AB) paved the way for characterizing attention as a two stage process (Chun & Potter, 1995). The first stage is a fast identification process in which targets are identified based on their features, such as colour or shape. The second stage is a slower, capacity limited process in which the identified target is consolidated in working memory for later report. When T2 arrives while T1 is still being consolidated, T2 can be identified but its consolidation is impaired, producing the AB. Supporting this theory, Vogel et al. (1998) found that ERP components associated with early attention and semantic processing (P1, N1, and N400) were present for T2 during AB trials, whereas the P300, associated with working-memory updating was supressed. Thus, the AB appears to reflect a limitation in consolidating T2 into working memory despite its initial identification. Other models have since proposed alternative mechanisms for the AB (Olivers & Meeter, 2008; Wyble et al., 2011).

Another model proposed to explain the AB is the episodic Simultaneous Type Serial Token (eSTST) model (Wyble et al., 2011). According to this model, attention is not allocated continuously over time but organised into discrete, temporally bounded attentional episodes. The onset and duration of these episodes depend on task demands and on the temporal relationship between successive targets. An attentional episode represents a period during which attention remains engaged, allowing one or more relevant targets to be encoded. In RSVP paradigms, these episodes operate over a relatively brief temporal window of approximately 150 ms. When two targets are presented in close temporal succession, such that the second target arrives before the first target’s processing is complete, the ongoing attentional episode is occupied with processing the first target, limiting the availability of attention for the second. This temporal limitation can contribute to the reduced detection of T2 during the AB. This sequence of transient processing episodes that unfold over a timescale of a few hundred milliseconds is theorised to be the underlying cause of AB.

One potential measure to track the dynamics of attention during an AB task is pupil response. Pupil response provides a non-invasive readout of cognitive processes such as attention and has been linked to activity in the neuromodulatory locus coeruleus–norepinephrine (LC–NE) system (Aston-Jones & Cohen, 2005; Joshi et al., 2016). Task-evoked pupil dilation has been widely used as an index of cognitive load, mental effort, and attentional engagement (Hess & Polt, 1964; Kahneman & Beatty, 1966; Mathôt, 2018; Ruuskanen & Mathôt, 2026; van der Wel & van Steenbergen, 2018). However, most conventional task-evoked pupillometry paradigms examine responses to a single target or to multiple stimuli sufficiently separated in time for the pupil response to develop as a relatively distinct response (Kafkas, 2024; van Rijn et al., 2012). In such paradigms, the magnitude of the dilation can be related to the cognitive demand associated with the target or task. However, the relatively slow temporal response of the pupil makes it difficult to distinguish responses to target stimuli that occur in rapid succession (as in RSVP), because the pupil response to one event may still be developing when the next occurs. As pupil responses evolve much more slowly than the underlying neural processes, responses to successive stimuli overlap extensively, making it difficult to associate changes in pupil size with individual attentional episodes with conventional analyses. So, a crucial question is, as pupil responses are driven by attention, can they be used to derive the associated latent attentional episode(s) responsible for target processing when targets are presented in rapid succession?

Hoeks & Levelt (1993) empirically showed that pupil response can be approximated with a canonical impulse response function (the ‘Pupillary Response Function’ or PRF). Building on this, Wierda et al. (2012) introduced a method to estimate latent attentional episodes (or, as they call it, “pulse”) from pupil responses to stimuli presented in rapid succession by deconvolving the pupil responses with the PRF. Deconvolution reverses the usual forward process. Instead of asking “What pupil response would these attentional events produce?”, deconvolution asks, “What pattern of attentional events could have produced the observed pupil response?” This makes it possible to estimate the temporal evolution of attentional engagement even when the corresponding pupil responses overlap substantially in time. This approach is conceptually analogous to hemodynamic response function (HRF) deconvolution in fMRI. Just as HRF deconvolution estimates the underlying neural events from overlapping BOLD responses (Glover, 1999), pupil deconvolution estimates the underlying attentional responses from overlapping pupil responses. However, this method of pupil deconvolution was primarily developed to model the pupil dilation that follows moments of increased attentional engagement (Wierda et al., 2012). This represents a critical limitation for visual paradigms, in which the pupil exhibits a biphasic response consisting of an early constriction phase reflecting the pupillary light reflex (PLR), followed by a slower dilation phase (Barbur et al., 2004; Korn & Bach, 2016). Therefore, to be generally applicable, the Pupillary Response Function (PRF) should reflect both constriction and dilation.Therefore, in the present study, we extended the existing pupil deconvolution model to incorporate both constriction and dilation phases through a dual-kernel model. This extension can handle the biphasic nature of the pupil response to a visual stimulus, which cannot be dissociated with the conventional single kernel approach. We apply this model to pupil data collected during an RSVP task using emotional faces as targets. The aim of the project was twofold. First, to examine whether the latent attentional episodes estimated from a biphasic pupil response reflect the dynamics of attention. As a more sophisticated test of our method, we focused on the valence based effects reported by Ray et al., (2020). If differences in AB reflect differential attentional demands for T1 and T2 across emotional valence, these differences should also be reflected in pupil responses or the amplitude of the attentional episodes derived through deconvolution. We therefore examined whether pupil responses and the resulting attentional episodes varied as a function of T1 and T2 valence.

Our model-derived attentional response captures key characteristics reported in prior literature, including temporal constraints in attentional allocation and the episodic nature of attentional deployment, consistent with established theories of the AB.

## 3. Method

A conventional power analysis to calculate the sample size was not feasible due to insufficient reporting of effect sizes and variance estimates in available literature. Instead, the sample size was calculated by comparing studies examining cognition driven pupil response and AB effect (Ray et al., 2020; Vilotijević & Mathôt, 2025; Wierda et al., 2012), in which sample sizes typically range from 25 to 40 participants. For our experiment, 37 participants were recruited from the University of Groningen. The participants received course credit for their participation. Following preprocessing and the application of predefined inclusion and exclusion criteria (see Section 3.3), data from 32 participants were retained for analysis (21 females, 11 males; age range: 17–24 years).

The study was exempted from full ethical review in accordance with the guidelines of the Ethical Committee of the Faculty of Behavioural and Social Sciences (EC-BSS) at the University of Groningen (study code PSY-2526-S-0083).

### 3.1 Task Paradigm: Attentional Blink with Emotional Faces

The task paradigm was adapted from Ray et al. (2020), who used emotional faces (Happy and Angry) as target stimuli and neutral faces as task-irrelevant distractors within an RSVP paradigm. In the present study, this paradigm was adopted because it has already been shown to elicit the AB effect and to measure both pupil constriction and dilation. Visual face stimuli elicit a pupillary light reflex in response to changes in visual input and also engage higher-level processes such as perception and attention. Images of emotional faces were selected from the Radboud Face Database (Langner et al., 2010). All images were pre-processed to remove non-essential facial regions (e.g., hair and clothing) using an oval mask and were luminance equalized to minimize low level visual confounds. Three emotional expressions were used: Happy (46 images), Angry (43 images), and Neutral (42 images) from the database.

Each trial started with a fixation point presented for 1000 ms at the centre of the monitor (Figure 1A), followed by rapid presentation of stimuli containing both target and distractor faces, alternating between face stimuli for 100 ms and a blank for 50 ms. All faces were presented at fixation, yet participants were instructed to maintain fixation and not make any eye movements during the RSVP stream. Each RSVP stream had 15 faces interlaced by a blank screen. In 20% of trials, only one emotional face was presented as the target (Figure 1B), and in the other 80% of trials, two emotional target faces were presented (Figure 1A). The target faces could be either happy or angry, with equal probability of them appearing. All the distractor faces were neutral. The first target (T1) could appear in serial positions 4, 5, or 6 relative to the stream’s onset (Figure 1C). So, the first target could appear as early as 550 ms in the RSVP stream, and the T2 can appear at serial positions 1, 2, 3, and 5 in the stream with respect to the T1 onset. This accounts for the stimulus asynchrony (SOA) of 150 ms (Lag 1), 300 ms (Lag 2), 450 ms (Lag 3) and 750 ms (Lag 5). All the lags were randomized within the experiment, and they also appeared an equal number of times. The trials with two targets can have one of the four possible pairs: Happy in T1 followed by Happy at T2 (HH), Happy at T1 followed by Angry at T2 (HA), Angry at T1 followed by Angry at T2 (AA) or Angry at T1 followed by Happy at T2 (AH). For the single target presentation, the target can be either Happy (H) or Angry (A). Following each RSVP stream, participants responded to a series of questions using keypress. First, they reported the emotion of the first target (Happy or Angry). Next, they indicated whether a second emotional target was present (“Yes” or “No”). If a second target was reported “Yes”, participants were prompted to indicate its emotional expression (Happy or Angry).

**Figure 1.**
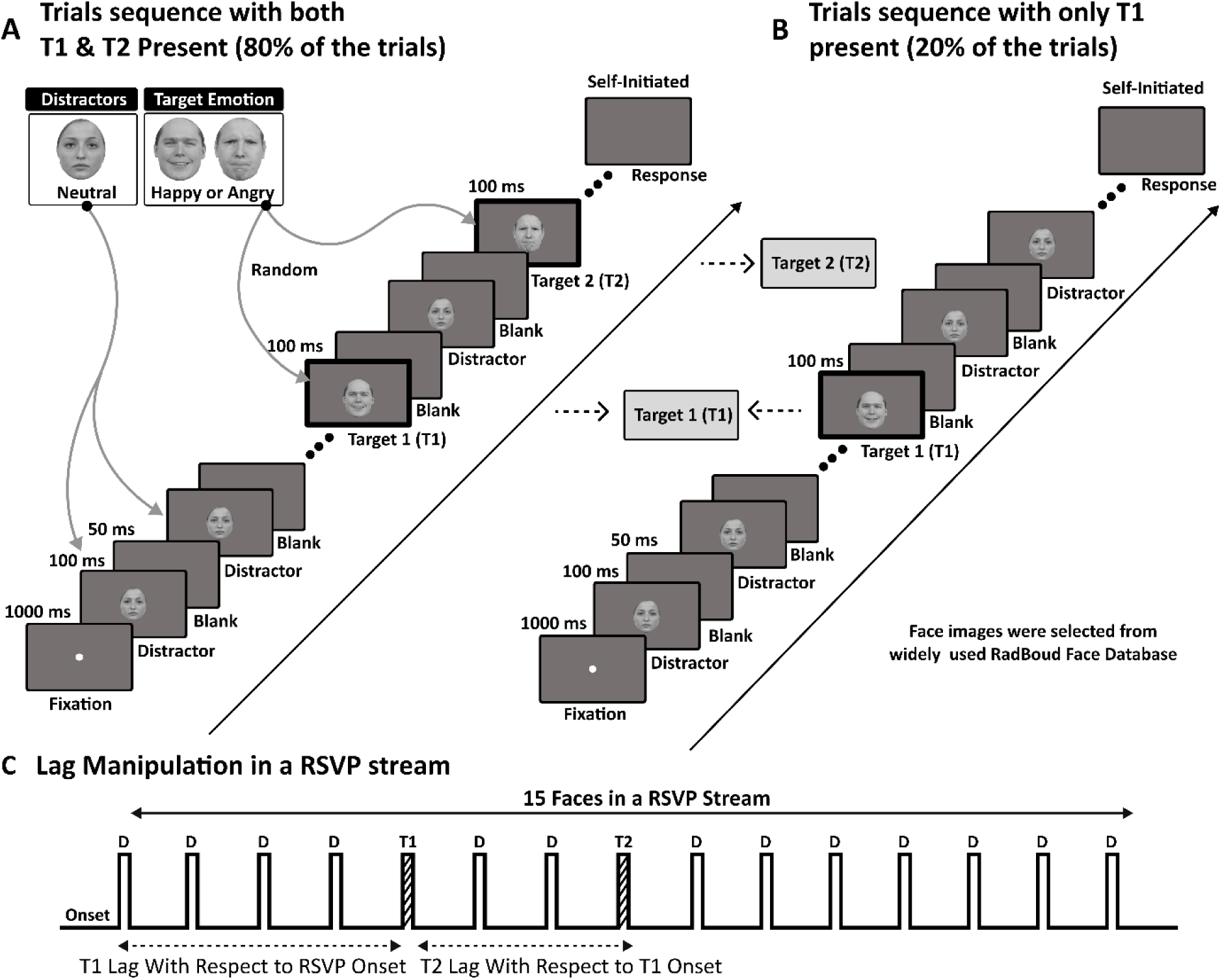
Attentional Blink Paradigm with Emotional Faces. (A & B) Each trial began with a central fixation period (1000 ms), followed by an RSVP stream of face stimuli. Stimuli were presented sequentially for 100 ms, interleaved with a 50ms blank interval. The RSVP stream consisted of 15 face stimuli, including neutral distractors and one or two emotional target faces (Happy or Angry). In 80% of trials, there will be two emotional targets; in 20%, there will be just one. (C) The temporal lag between T1(first target) and T2 (second target) was manipulated across trials, corresponding to stimulus onset asynchronies of 150 ms (Lag 1), 300 ms (Lag 2), 450 ms (Lag 3), and 750 ms (Lag 5). D represents the distractors.

### 3.2 Experimental Setup and Procedure

Participants were provided with instructions about the experiment. Upon going through the details, they provided written consent to participate in the experiment. The study was performed in a dimly lit room, with a mean light luminance across all the sessions of 38 lux. The experiment was designed in OpenSesame (Mathôt et al., 2012) using the PyGaze Python package for eye tracking and was presented on a 27-inch PROLITE G2773HBS-GB1 (EOL) monitor with a resolution of 1920×1080 and a refresh rate of 60 Hz. Pupil data were recorded using an EyeLink 1000 at a sampling rate of 500 Hz (SR Research). Pupil size was recorded monocularly (right eye) and measured in arbitrary units of pupil area. The participants were seated 60 cm from the computer monitor. Throughout the experiment, the participant’s head and chin were supported by a chinrest to minimise head movement.

The experiment started with an eye tracking calibration-validation procedure using 9 on screen marker points, followed by an instruction on the screen. Once initialized, the instruction was followed by a practice block consisting of 20 trials. A single point drift correction procedure was initiated before every trial to bring the eye to the centre of the monitor. The main experiment began upon completion of the practice block and was divided into two parts. Each part consisted of 120 trials, followed by a five minute break.

### 3.3 Pupil Data Preprocessing

Participants were included in the analysis only if their accuracy in detecting Target 1 (T1) exceeded 70%, ensuring only those sufficiently engaged with the task were retained (Ray et al., 2020). Of the 37 participants recruited, 32 met this criterion and were included in subsequent preprocessing and analysis.

Pupil data were pre-processed using custom Python scripts following established guidelines for pupil data analysis (Mathôt & Vilotijević, 2022). Continuous pupil recordings were first segmented into fixation and RSVP epochs for each trial. Eye blinks were identified based on event markers provided by the eye tracker, and blink data were reconstructed using cubic spline interpolation using functions available in the DataMatrix package. The reconstructed pupil signal was down sampled from 500 Hz to 100 Hz. Baseline correction was performed by subtracting the mean pupil size computed over the final 100 ms of the fixation period preceding stimulus onset from the entire trial.

### 3.4 Method to Infer latent attentional episodes from pupil response

To estimate the latent attentional episodes from the pupil response, we used a model based deconvolution approach adapted from the work of Hoeks & Levelt, (1993) and Wierda et al., (2012). Wierda et al., (2012) demonstrated that the relatively slow pupil response can be modelled as the linear convolution of a series of latent attentional episodes with a canonical pupil impulse response function (PRF), thereby providing an estimate of latent attention at substantially higher time resolution than the raw pupil signal response. In the present study, repeated presentation of emotional face stimuli elicited pupil constriction followed by dilation. To accommodate these additional physiological dynamics, we adapted the deconvolution framework by modelling the observed pupil signal as the combined contribution of both constriction and dilation phases (Figure 2).

**Figure 2:**
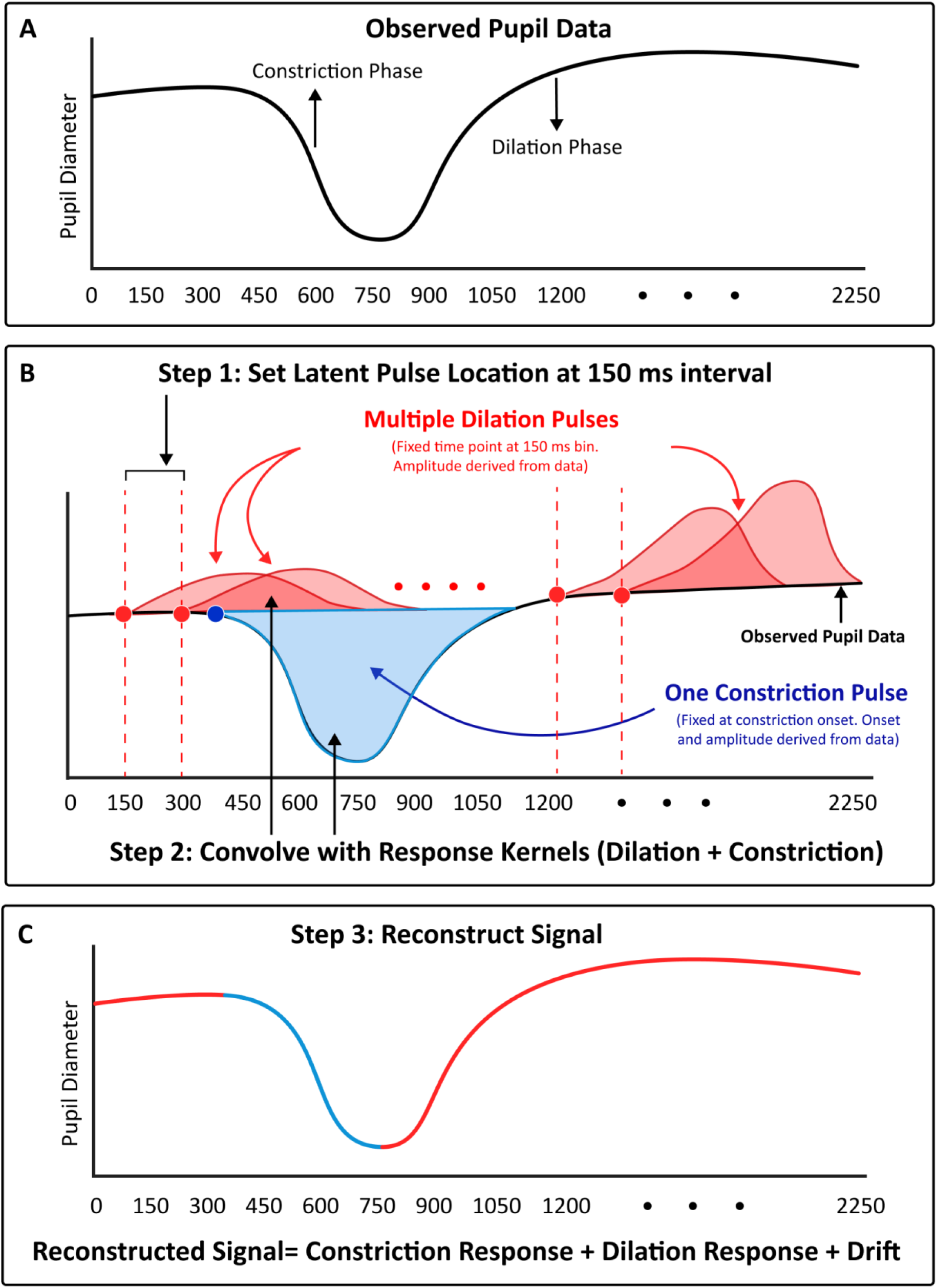
Schematic overview of the dual-kernel pupil deconvolution approach used to estimate latent attentional response from pupil response. (A) Representative pupil response during the RSVP task, showing the combined effect of stimulus evoked constriction and task related dilation responses. (B) The dual-kernel model consists of a constriction kernel (blue) and dilation kernels (red). The constriction component is anchored to the estimated PLR onset, whereas dilation kernels are placed at successive stimulus onsets (150 ms apart), with their amplitudes estimated from the data. The constriction kernel is convolved with one point at constriction onset, while the dilation kernel is convolved multiple times at a 150 ms bin (attentional pulse location). (C) Reconstructed pupil response obtained by combining the constriction and dilation components.

The core assumption of our model is that the phases of pupil dynamics can be approximated as a linear time invariant (LTI) system, which means that the strength or shape of a pupil response at one moment does not depend on the pupil response at another moment. This assumption is supported by empirical evidence showing that pupil responses to brief stimuli exhibit a biphasic temporal (Figure 2A) response that is invariant across conditions, when measured under controlled luminance conditions (Korn & Bach, 2016). We do not claim that the pupil system is strictly linear or time invariant in a physiological sense. Rather, the LTI formulation is used as a computational approximation adopted in prior pupillometry work (Cai et al., 2023).

#### 3.4.1 Impulse Response Functions of the Pupil

The transformation from latent events to observed pupil responses was modelled using impulse response functions (Pupil Response Functions; PRF) that characterise the pupil response following a visual stimulus. Hoeks & Levelt. (1993) characterized the pupil response as the output of a system driven by neural or cognitive events, where each event evokes a temporally extended response with a gradual rise and slow decay (see representative schematic in Figure 2B). They characterised the PRF using an Erlang-Gamma function to approximate the temporal profile and experimentally demonstrated that this function best explains the pupil’s impulse response. Extending this model, Wierda et al. (2012) introduced a model based deconvolution approach to estimate latent attentional episodes from pupil responses in a paradigm involving fast paced stimulus presentation. In this approach, the pupil signal is thought to arise from attentional episodes that emerge at high temporal resolution. In their method, the pupil dilation component was modelled using the PRF proposed by Hoeks & Levelt (1993). The dilation kernel was defined as:

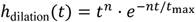

where *t* denotes time since event onset, *n* controls the shape of the response (the steepness of the rise), and *t*_max_ determines the time to peak. In the present study, we used parameter values *n* = 10.1 and *t*_max_ = 930 ms, which produce a delayed peak and prolonged decay consistent with empirically observed and reported parameters Hoeks & Levelt, (1993).

While the method proposed by Wierda et al., (2012) models the dilation part, it does not account for the early constriction phase elicited by any visual stimuli. Physiologically, pupil constriction is mediated by parasympathetic pathways and exhibits a faster onset, earlier peak, and shorter duration compared to dilation (Korn & Bach, 2016). A target stimulus may occur during the early constriction phase, which a dilation only model cannot adequately capture. As a result, the response to that stimulus may be underestimated. To capture this constriction phase, we introduced a second kernel that represents the constriction component of the pupil response. The constriction kernel was derived from the same parametric family as the dilation kernel but parameterized to reflect faster temporal dynamics and inverted polarity. It was defined as:

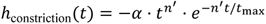

where *α* scales the amplitude of the constriction response, *n*^′^controls the shape of the kernel, and *t*_max_ which is the same as the dilation kernel (930 ms), determines the time to peak response. The negative sign indicates a decrease in pupil diameter. The constriction kernel was defined using *n*^′^ = 6 and *α* = 0.6. The value of *α* was chosen arbitrarily to scale the constriction response relative to the dilation response, whereas *n*^′^determine its temporal shape. The shape parameter *n*^′^, produces a faster rise and earlier peak than the dilation kernel (*n*), consistent with empirical observations of PLR response (Korn & Bach, 2016). The temporal position of the constriction response was determined separately by estimating the PLR latency from the steepest negative pupil slope within the first 400 ms after stimulus onset, as pupil constriction happens within 350-400 ms after stimulus presentation (Joshi et al., 2016). The constriction kernel was then anchored at this latency (Figure 2B). The kernels were normalized before model fitting, and their amplitudes were estimated from the data during optimization. The kernels define only the temporal transformation of latent events, whereas their amplitudes are estimated during model fitting. As such, the inferred attentional episodes are driven primarily by data driven estimation of amplitude rather than by the specific choice of kernel parameters.

#### 3.4.2 Representation of Latent Events

Each stimulus in the RSVP stream corresponded to one latent attentional event. This discretization does not assume that attentional events occur at every predefined time point. Rather, it provides a structured set of candidate temporal locations from which the model infers when attentional allocation most likely occurred. At each time point, the amplitude of the corresponding episode was treated as a free parameter. The latent attentional input can thus be represented as a discrete time sequence *i*_attn_(*t*), defined over the temporal grid, where each element corresponds to the amplitude of an episode at time *t*. These amplitudes represent the strength of the underlying latent event and are estimated during model fitting. While the temporal locations of episodes were fixed, their amplitudes were not predetermined. Instead, they were inferred from the data by optimizing the fit between the model predicted and observed pupil response. As a result, the model determines which time points contribute to the observed response by assigning higher amplitudes to those events and lower amplitudes to others. The resulting amplitudes should be interpreted as model derived estimates of underlying latent neural or cognitive events that give rise to the observed pupil signal, rather than as direct measures.

#### 3.4.3 Modelling of the Pupil Signal

Given the impulse response functions and the representation of latent events described above, the observed pupil signal was modelled as the output of a process that transforms discrete sensory and attentional events into continuous pupil responses. Each latent attentional event was convolved with the dilation kernel. This reflects the assumption that each stimulus triggers an attentional process. In contrast, only a single time point was convolved with the constriction kernel; this reflects the assumption that the onset of the RSVP stream effectively triggers a single constriction event. In addition, a linear drift component was included to account for slow baseline fluctuations unrelated to task events. The predicted pupil signal 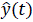 can be expressed as:

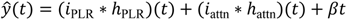

where *i*_PLR_(*t*) and *i*_attn_(*t*) denote the latent sensory and attentional episode sequences, ℎ_PLR_(*t*) and ℎ_attn_(*t*) are the corresponding constriction and dilation kernels, ∗ denotes a convolution operation, and *β* is the linear drift parameter. The predicted pupil signal is thus constructed as the sum of these components (Figure 2C).

#### 3.4.4 Parameter Estimation and Optimization

Free model parameters were estimated by fitting the model to each participant’s average pupil response. The amplitude of attentional episodes and the linear drift parameters were treated as free parameters and optimized to minimize the mean squared error between the predicted and recorded pupil responses, defined as 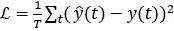. Optimisation was performed using a bounded quasi-Newton method (L-BFGS-B), as it provides efficient estimation under bounded constraints (Liu & Nocedal, 1989). Amplitudes were constrained to be non negative because they would lead to an additive contribution to the attentional episodes (optimization method adopted from Wierda et al., 2012), while the drift parameter was allowed to vary within a range around zero. To mitigate local minima convergence, the optimization procedure was repeated fifty times with randomized initializations, and final parameter estimates were obtained by averaging across the fifty iterations.

#### 3.4.5 Model Output and Derived Measures

The model’s primary output was the estimated amplitude of latent attentional episodes over time. The attentional episodes, defined on the fixed temporal grid, were interpreted as a continuous estimate of how attention evolved over time, with higher amplitude indicating stronger attentional engagement at corresponding time points. The drift parameter was not analyzed.

#### 3.4.6 Implementation Details: *PupilDeconvolve* Package

To facilitate reproducibility and application of the proposed framework, we developed *PupilDeconvolve*, an open-source Python package implementing the complete dual kernel deconvolution pipeline described in this study. All modelling and optimization procedures were implemented in Python (version 3.14) using standard scientific computing libraries (Numpy, Pandas, SciPy). Convolution operations are performed using Fast Fourier Transform (FFT) based convolution (fftconvolve) implemented in SciPy, providing efficient evaluation of the forward model during iterative optimization.

This deconvolution method and package are applicable to experimental paradigms involving the rapid presentation of stimuli in which biphasic pupil response to multiple target stimuli overlap substantially, including RSVP, visual search, and other time critical cognitive tasks where conventional analyses of pupil responses are limited by the slow temporal dynamics of the pupil response. The package can be installed using ‘pip install pupildeconvolve’. Source code, documentation, tutorials, and example datasets are available at https://github.com/Sangramjit/pupildeconvolve

#### 3.4.7 Running the Deconvolution (*pupildeconvolve*) Package

The *pupildeconvolve* package requires a dataframe containing pre-processed pupil time series data, with each row corresponding to a participant and the pupil time series provided as separate columns. The user must specify the pupil data columns, the participant identifier, the sampling rate, and either the interval between successive attentional events or the explicit timing of those events. For each participant, PLR latency is estimated automatically from the pupil signal. The estimated PLR response is then incorporated into the subsequent deconvolution. The process will return a dataframe containing participant level deconvolution estimates. The output includes the participant identifier, the estimated slow baseline drift (slope), PLR latency and the estimated amplitude of each attentional episode.

## 4. Data Analysis

Behavioural performance was analysed to verify the presence of the AB effect and to assess the influence of emotion on the AB magnitude. Analysis was focused on T2 report accuracy as a function of temporal lag and emotional pairing (emotion at T1 and T2). Trials were categorized based on the combination of T1 and T2 emotions (Happy–Happy, Happy–Angry, Angry–Happy, Angry–Angry) and lag (Lag 1, 2, 3, and 5). Accuracy data were analyzed using generalized linear mixed effects model (GLMMs), implemented in the lme4 package in R. For the two-target condition, fixed effects included Emotion pair and lag, as well as their interaction. Participant was included as a random effect with a random intercept for lag to account for individual differences in overall accuracy and AB across lags. When significant main effects or interactions were observed, post hoc comparisons were conducted using estimated marginal means with Tukey’s adjustment for multiple comparisons. For the single target condition, accuracy was also analysed using a GLMM with Emotion (Happy vs. Angry) as a fixed effect and Participant as a random intercept. Pupil and attentional episode time courses were analysed to see differences across conditions. To ensure sufficient statistical power, analyses were performed on condition wise aggregated data (e.g., No Blink vs. Blink) rather than separating trials by lag, as lag wise segmentation resulted in an insufficient number of trials per participant. The first and last attentional episodes were excluded from analysis, as they correspond to boundary conditions.

Statistical comparisons were conducted using a cluster based permutation test. For factorial analyses, repeated measures ANOVA was computed at each time point to assess main effects and interactions (Blink × Emotion), and cluster level statistics were derived from these tests. Significance was determined by comparing observed cluster statistics against a permutation based null distribution. Clusters were defined as contiguous time points exceeding a predefined statistical threshold (α = 0.05). For each cluster, a cluster-level statistic was computed as the sum of the test statistics within the cluster. Statistical significance was evaluated using a permutation procedure in which condition labels were randomly shuffled within participants. For each permutation, the maximum cluster level statistic was retained to construct a null distribution. This procedure was repeated 5,000 times and observed cluster statistics were compared against this null distribution to obtain cluster level p-values, thereby controlling for multiple comparisons.

## 5. Result

### 5.1 Behavioural analysis

To establish the presence of the AB and its modulation by emotion, we have analyzed the T2 report accuracy (Figure 3A) as a function of lag (1, 2, 3, 5) and emotion pair (AA, AH, HA, HH). GLMM analysis revealed a significant main effect of lag; χ²(3) = 235.17, p < .001, emotion pair; χ²(3) = 249.90, p < .001, as well as a significant emotion pair × lag interaction; χ²(9) = 126.10, p < .001. Post hoc comparisons showed that T2 accuracy at Lag 1 was significantly lower than at Lag 2 and Lag 3 across all emotion pair conditions (p < .001). At later lags, accuracy increased further from Lag 3 to Lag 5 for AA (z = −5.75, p < .001), AH (z = −4.66, p < .001), and HA (z = −5.62, p < .001), but not for HH (z = −2.00, p = .23). Detailed pairwise comparisons are provided in the Supplementary Result. Overall, T2 accuracy was significantly reduced at the shortest lag (Lag 1) and increased as the temporal separation between T1 and T2 increased. Across all emotional pair conditions, the consistent reduction in T2 detection accuracy at Lag 1 relative to later lags indicates the presence of AB. The significant Emotion Pair × Lag interaction further suggests that the magnitude of this blink varied as a function of the emotion of the target pair.

**Figure 3:**
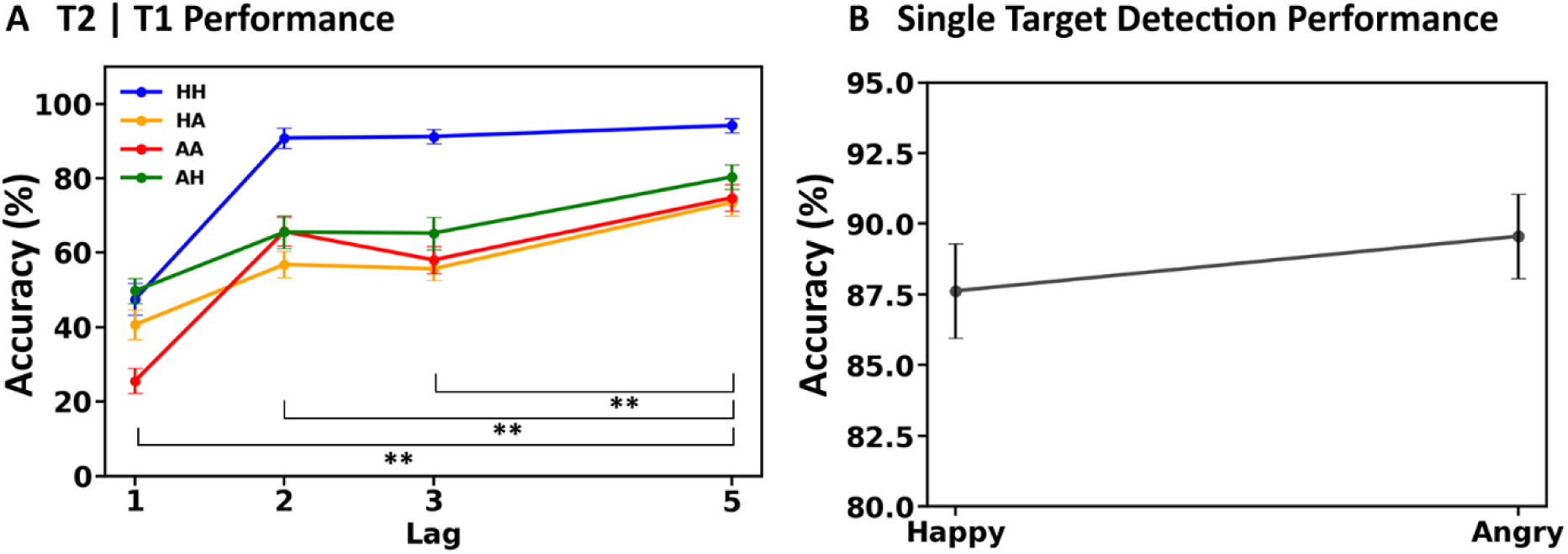
Behavioural performance in the RSVP task. (A) T2 report accuracy (%) as a function of lag (Lag 1, 2, 3, and 5) for each emotional pairing: Happy–Happy (HH), Happy–Angry (HA), Angry–Angry (AA), and Angry–Happy (AH). Accuracy is plotted as the mean across participants, with error bars representing the standard error of the mean (SEM). A clear attentional blink effect was observed, with reduced T2 accuracy at short lags (Lag 1) relative to longer lags (Lag 3 and Lag 5) across all conditions. * Represents a significant difference. (B) Single-target detection accuracy (%) for Happy and Angry targets. Error bars represent SEM.

In the single target condition (Figure 3B), accuracy did not differ between Happy and Angry targets χ² (1) = 1.91, p = .167.

### 5.2 Analysis of Pupil Response

The pupil response (as measured directly, without applying deconvolution) was analysed to examine whether different conditions influenced it. Trials were categorized into ‘No Blink’ trials, defined as trials in which both T1 and T2 were correctly reported and ‘Blink’ trials, defined as trials in which T1 was correctly reported but T2 was missed/ incorrectly reported. Pupil responses were also compared between No Blink and single target trials. Statistical comparisons were conducted using cluster-based permutation tests as described in the ‘Data Analysis’ section. Cluster-based permutation tests combined with repeated measures ANOVA to assess effects [Blink Effect (Blink and No Blink) × Emotion (HH, HA, AA, AH)]. Across all comparisons, no significant difference was observed at any time point (Figure 4A, C, E, G) for any of the conditions (e.g., No-Blink vs Blink). In the present RSVP paradigm, multiple stimuli (15 items) were presented within a short time window (∼2250 ms), leading to substantial overlap of pupil responses evoked by successive stimuli. Given the slow rise and prolonged decay of pupil dilation, responses to individual events accumulate and cannot be temporally dissociated in the observed signal. As a result, it is difficult to directly observe differences in attentional processing from the pupil response alone. Accordingly, subsequent analyses focus on the analysis of the latent attentional episodes recovered through model based deconvolution of the pupil signal as explained in section 3.4 of the ‘Method’ section.

**Figure 4:**
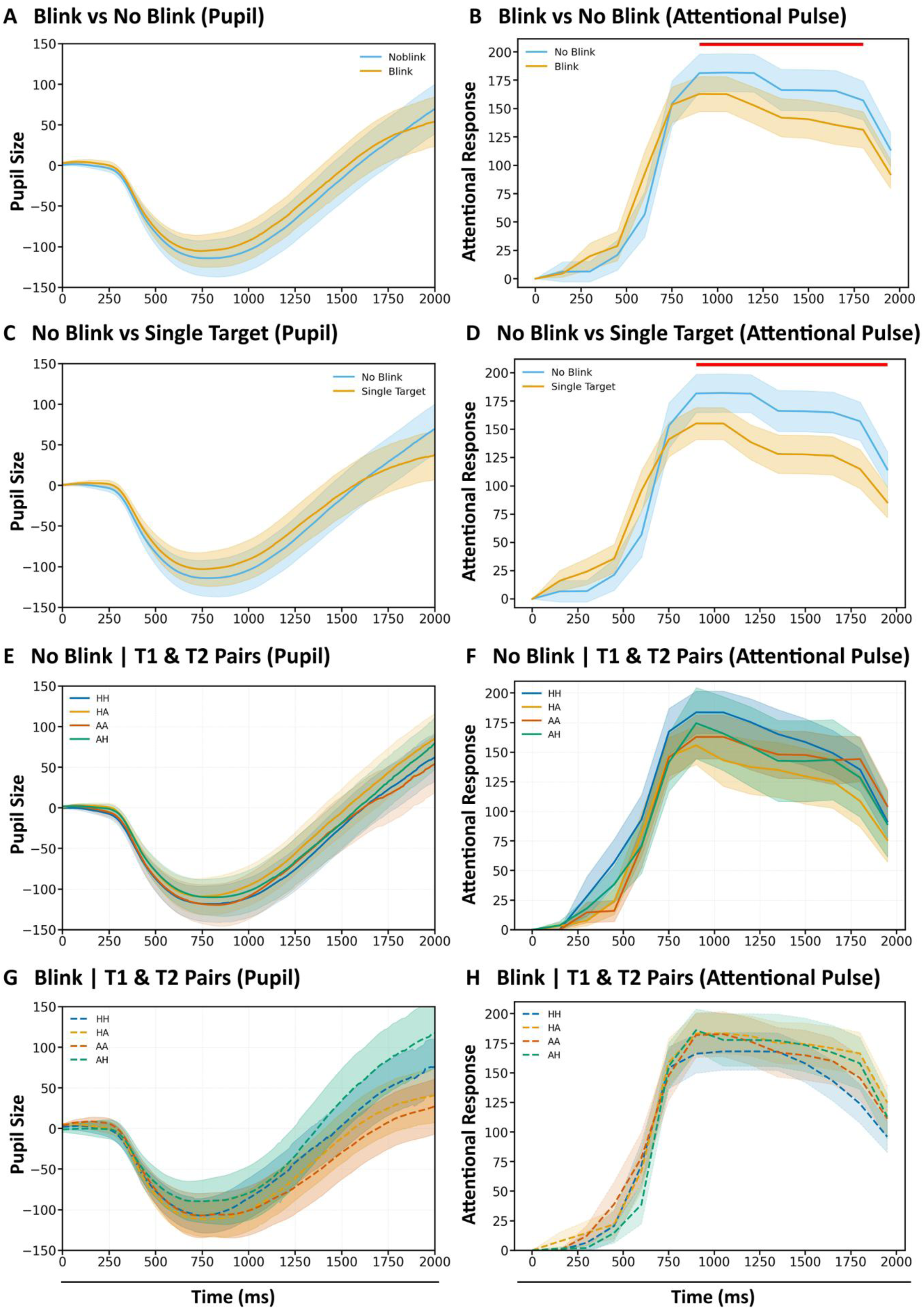
Effects of target conditions on pupil response and inferred attentional episodes. Panels show time courses (0–2000 ms) of mean pupil size (left column) and model derived attentional episodes amplitude (right column), with shaded areas indicating ± SEM. The red bar reflects the significant cluster. The plots were from the onset of the RSVP stream (0 ms). (A–B) Comparison of Blink vs No Blink conditions. (C–D) Comparison of No Blink vs Single Target trials. (E–F) No-blink trials split by T1–T2 pair types (HH, HA, AA, AH). (G–H) Blink trials split by T1–T2 pair types.

### 5.3. Latent attentional episodes differ among conditions

Next, we compared the amplitude of the attentional episodes (derived from pupil response with deconvolution) between No-Blink and Blink. As shown in Figure 4B, the amplitude in the No-Blink condition (blue line) is consistently higher than in the Blink condition (orange line). Mean amplitude was higher in the No-Blink condition (Mean:111.26, SEM:11.07) than in the Blink condition (Mean:101.37, SEM:9.82). A cluster-based permutation test revealed a significant cluster between 900 and 1800 ms, indicating that the two conditions differ in amplitude within this time window (p < .001, cluster mass = 18.61). We next compared amplitudes between the single target and No-Blink conditions. The amplitudes differed significantly (Figure 4D). Mean amplitude was higher in the No-Blink (blue line) than in the single target (orange line) condition (Mean:96.06, SEM:9.35). A cluster-based permutation test identified a significant cluster between 900 and 1950 ms (p < .0001, cluster mass = 28.05).

Finally, we examined whether the amplitude of attentional episodes differed across emotional conditions. Behaviourally, emotion influenced detection accuracy, with higher performance for HH pairs than for AA (Figure 3A), particularly at early lag (Lag 1). To assess whether these behavioural differences were reflected in amplitude, we compared Blink and No-Blink trials within each emotion pair. Due to limited trial counts, trials were aggregated across lags within each condition. As shown in Figure 4F (No-Blink) and Figure 4H (Blink), attentional episode time courses for different emotion pairs, HH (blue), HA (orange), AA (red), and AH (green), overlap. Statistical comparison revealed no significant interactions (Blink × Emotion; all cluster-based permutation tests were non-significant) or main effects of Emotion and Blink. Although emotion modulated behavioural performance, these differences were not reflected in the amplitude of the attentional episode.

## 6. Discussion

In this study, we examined how emotional targets presented in close temporal succession influence the deployment of attention and whether these differences are reflected in the pupil response or in the latent attentional episodes estimated from the biphasic pupil response. Specifically, we addressed two questions. First, we examined whether latent attentional episodes estimated from the slow, biphasic pupil response to emotional targets reflect the temporal limitations of attention. Second, we investigated whether the emotional valence of targets presented in close temporal succession modulates the magnitude of these attentional episodes.

We examined how emotional targets influenced AB. In the behavioural data, T1 accuracy remained high across emotional conditions, whereas T2 accuracy showed a clear AB effect, particularly at short lag (Lag 1) (Figure 3A & B). We also found that emotional targets modulate the AB effect. This pattern is consistent with findings from Ray et al. (2020).

Next, we examined pupil responses during the RSVP task. Pupil responses are closely linked to activity in the locus coeruleus–noradrenergic (LC–NE) system, which plays a central role in regulating arousal (Joshi et al., 2016) and in mediating the temporal capacity of attention (Nieuwenhuis et al., 2007). However, pupil response is substantially slower than the LC-mediated response, which peaks around 100-150 ms following target presentation (Aston-Jones et al., 1994; Clayton et al., 2004), whereas the pupil starts responding around 300-450 ms after stimulus onset (Joshi et al., 2016). Thus, when stimuli (targets and distractors) are presented in rapid succession, their pupil responses overlap substantially, and the measured response reflects the cumulative effect of multiple stimuli rather than a distinct response to each target. This is evident in Figures 4A, C, E, and G, in which pupil response marked by a constriction followed by a dilation phase is observed for a time duration (2000 ms after RSVP onset) over which RSVP was presented. A single response represents the cumulative response to stimuli presented within RSVP and cannot reliably distinguish responses to individual events/targets presented in rapid succession, limiting its ability to capture the temporal dynamics of attention. This limitation extends beyond RSVP paradigms to other paradigms in which target stimuli were presented in close succession, such as naturalistic viewing and sequential working memory tasks (Camenzind et al., 2026; van Rijn et al., 2012).

Crucially, to address this limitation of pupil response, we extended the model proposed by Hoeks & Levelt, (1993) and later used by Wierda et al., (2012) to estimate latent attentional episodes from biphasic pupil response via deconvolution. The method proposed by Wierda et al., (2012) has primarily modelled the dilation part of the pupil response, assuming that attentional events occur during this phase. However, the pupil response to visual stimulation is biphasic, consisting of an early reflexive constriction followed by a slower dilation phase (Korn & Bach, 2016). This is particularly important in the RSVP paradigm because T1 can occur at different points within the pupil response, including during the early constriction phase or later dilation. Therefore, we developed a dual-kernel deconvolution model that separately captures the constriction and dilation components of the pupil response (Figure 2).

(Wierda et al., 2012) observed two distinct groups of attentional episodes, with higher amplitudes than neighbouring points: the first around 500 ms and the second around 1000 ms, for the condition in which both targets were detected and reported properly (No-Blink). The first episode was temporally locked to T1 onset, whereas the second episode occurred at the expected T2 time point. This temporal correspondence led them to interpret the two episodes as reflecting the processing of T1 and T2, respectively. In our data, a peak was defined as a local maximum in the attentional response. Consecutive time points with similar elevated amplitudes were grouped into a single peak. A new peak was identified only when the response decreased between two elevated regions, forming a distinct local valley. Our estimated latent attentional episodes revealed two broad, distinct groups when both targets were correctly detected (No-Blink; Figure 4B, Blue Line). The first group was identified within 700–900 ms (the time window of our T1 presentation), and the second within 1100–1400 ms (their latency differ significantly; p<0.001). Two distinct groups were also observed in the Blink condition, in which T1 was detected but T2 was missed. However, the magnitude was significantly lower than in the No-Blink condition (Blink; Figure 4B Orange Line). The No-Blink condition also showed a consistently greater attentional magnitude than both the Blink (Figure 4B) and single-target (Figure 4D) conditions, suggesting greater attentional engagement when two targets were successfully processed. This pattern is also consistent with findings from Wierda et al. (2012), who reported greater attentional magnitude when both targets were correctly reported (No-Blink) than when T2 was missed (Blink). Evidence from fMRI studies has similarly shown that the frontoparietal network, a brain network associated with attentional control and target selection (Corbetta & Shulman, 2002), shows reduced activity when T2 is not successfully reported (Marcantoni et al., 2003; Marois et al., 2004). ERP studies have shown that missed T2 reporting is associated with reduced activity around 200 ms after stimulus onset, which is associated with attentional selection (Luck et al., 1996; Martens et al., 2006). This substantiates the significant decrease in amplitude of attentional pulses in the Blink condition as compared to the No-Blink condition.

Interestingly, we also observed two distinct groups of attentional episodes in the single target condition, although their magnitudes were lower than those in the No-Blink condition (Figure 4D). Wierda et al. (2012) observed a very shallow increase in the second group of attentional episodes in their single target condition, which they interpreted as a possible expectancy effect associated with T2. In our data, the single target condition showed a distinct second attentional response at approximately 1100–1300 ms. This may reflect the expectancy effect, arising from the relatively low probability of single target trials. In our study, participants encountered a single target in only 20% of trials (Figure 1B), whereas 80% of trials contained two targets, and Wierda et al. (2012) has 18% of trials with a single target. Thus, following the first target, participants may have maintained an expectation of a second target, producing a later attentional response even when no second target was presented. Also, the relatively stronger amplitude corresponding to the expectancy effect observed in our data, compared with that reported by Wierda et al. (2012), may be attributable to the valence and arousal of the target emotional faces, which differ from those of the letter targets used by Wierda et al. (2012). This aligns with a study that showed expectancy enhanced neural responses to emotional, but not neutral pictures (Bermpohl et al., 2006).

We also compared the magnitude of attentional episodes within the emotion pairs presented in T1 and T2. Although the behavioural results showed that emotion pairs presented at T1 and T2 influenced the magnitude of the AB, with happy T2 targets consistently producing higher accuracy, we found no significant differences in the magnitude of the attentional episodes across emotion pairs (Figure 4F, H). There are two possible explanations for this. First, emotional effects were primarily observed at lags, as shown in the behavioural result (Figure 3A). Therefore, combining trials across lags may have obscured emotion specific differences. However, analysing each emotion pair separately at each lag would substantially reduce the number of trials, especially at Lag 1, making the deconvolution estimates noisier and less reliable. Second, emotional valence and arousal have been shown to influence the temporal dynamics of the pupillary response in a detection task in which the aim was to detect only one stimulus (no rapid presentation of distractors) (Astudillo et al., 2018). In our RSVP paradigm, 15 stimuli (target + distractors) were presented within an approximately 2100 ms time window, while the pupil response to a single emotional stimulus can unfold over nearly 2000 ms. Thus, responses to multiple emotional targets are likely to require a longer time window to reflect or even influence the pupil response, which can’t be resolved with our RSVP time window. These methodological constraints may have limited the ability to detect emotion specific influence in the estimated attentional episodes.

Taken together, the present findings demonstrate that the dual-kernel deconvolution approach provides a method for analysing the dynamics of latent attentional episodes from the biphasic pupil response, which was not possible with any previous method. The characteristics of the estimated attentional episodes were consistent with theories of the Attentional Blink, as evidenced by differences in attentional magnitude between the No-Blink and Blink conditions and between the No-Blink and single-target conditions. The attentional episodes also align with the episodic nature of attentional deployment (Wyble et al., 2011), in which we found distinct groups of episodes that may be related to the processing of T1 and T2. Importantly, this method offers a non-invasive measure of attentional dynamics that can complement neuroimaging methods such as EEG and MEG. Future work should focus on establishing the correspondence between model derived response and neural markers of temporal capacity of attention, as well as extending the approach to more complex and naturalistic tasks.

## 7. Limitations of the Study and Computational Modelling

Despite its utility in estimating latent attentional dynamics from pupil signals in rapid visual paradigms, the present approach has several methodological and modelling limitations. First, the number of trials per condition constrained the granularity of analysis. Dividing data across lag, emotion, and condition resulted in insufficient observations per condition, necessitating aggregation across lags for pupil and attentional episodes analyses. While this improves statistical reliability, it limits the ability to resolve lag specific attentional episodes. Second, the model assumes that pupil responses can be approximated as a linear time-invariant (LTI) system. Although widely adopted in pupillometry, the physiological pupil system exhibits nonlinear and state dependent properties, and thus the estimated attentional episodes should be interpreted as approximations rather than direct measurements. Finally, the model requires iterative optimization over a high dimensional parameter space, with each episode’s amplitude treated as a free parameter. Although computational efficiency is improved through FFT based convolution and constrained optimization, the approach remains computationally demanding and may limit scalability and real time use cases.

## Data Availability

The dataset is publicly available through Zenodo: 10.5281/zenodo.23056404

Analysis code is available through GitHub: https://github.com/Sangramjit/Model-Based-Pupil-Deconvolution

## Author Contribution

Sangramjit Maity: Conceptualization, Methodology, Data curation, Investigation, Project administration, Validation, Visualization, Writing – original draft.

Sebastiaan Mathôt: Methodology, Investigation, Resources, Project administration, Supervision, Validation, Writing – review and editing.

Sonia Baloni Ray: Conceptualization, Methodology, Resources, Project administration, Supervision, Writing – review and editing.

## Supporting information

Supplemental Analysis 1

