## Supplemental Analysis 1 for "Model-based pupil deconvolution reveals latent attentional dynamics"

**Supplementary:**

1. **Behavioural result: Post Hoc Comparisons of T2 Accuracy**

Following the significant emotion pair × lag interaction in the GLMM, post hoc pairwise comparisons were conducted to examine differences in T2 accuracy across lags within each emotional target-pair condition. For the AA condition, T2 accuracy at Lag 1 was significantly lower than at Lag 2 ($z = -9.23$, $p < .001$), Lag 3 ($z = -7.87$, $p < .001$), and Lag 5 ($z = -11.28$, $p < .001$). Accuracy at Lag 2 was not significantly different from Lag 3 ($z = -2.19$, $p = .17$) but was lower than at Lag 5 ($z = -3.09$, $p = .01$). Accuracy at Lag 3 was also significantly lower than at Lag 5 ($z = -5.75$, $p < .001$). For the AH condition, accuracy at Lag 1 was significantly lower than at Lag 2 ($z = -4.21$, $p < .001$), Lag 3 ($z = -3.81$, $p < .001$), and Lag 5 ($z = -8.27$, $p < .001$). Lag 2 and Lag 3 did not differ significantly ($z = -0.01$, $p = 1.00$), whereas accuracy at Lag 2 was lower than at Lag 5 ($z = -4.93$, $p < .001$). Accuracy at Lag 3 was also significantly lower than at Lag 5 ($z = -4.66$, $p < .001$). For the HA condition, accuracy at Lag 1 was significantly lower than at Lag 2 ($z = -4.59$, $p < .001$), Lag 3 ($z = -3.95$, $p < .001$), and Lag 5 ($z = -8.51$, $p < .001$). Accuracy at Lag 2 did not differ significantly from Lag 3 ($z = 0.40$, $p = 1.00$), but was significantly lower than at Lag 5 ($z = -5.29$, $p < .001$). Accuracy at Lag 3 was also significantly lower than at Lag 5 ($z = -5.62$, $p < .001$). For the HH condition, accuracy at Lag 1 was significantly lower than at Lag 2 ($z = -9.77$, $p < .001$), Lag 3 ($z = -13.49$, $p < .001$), and Lag 5 ($z = -10.08$, $p < .001$). No significant differences were observed between Lag 2 and Lag 3 ($z = 0.003$, $p = 1.00$), Lag 2 and Lag 5 ($z = -1.78$, $p = .44$), or Lag 3 and Lag 5 ($z = -2.00$, $p = .23$).

Overall, T2 accuracy was consistently reduced at Lag 1 compared with longer lags across all four emotional target pair conditions.
